# A novel resistance reversion mechanism in a vancomycin-variable *Enterococcus faecium* strain

**DOI:** 10.1101/2023.04.28.538149

**Authors:** Ross S. McInnes, Ann E. Snaith, Steven J. Dunn, Maria Papangeli, Katherine J. Hardy, Abid Hussain, Willem van Schaik

## Abstract

**Objectives:** To investigate an outbreak of *Enterococcus faecium* in a hospital haematology ward and uncover the mechanism of a vancomycin resistance phenotype-genotype disparity in an isolate from this outbreak.

**Methods:** Whole genome shotgun sequencing was used for the phylogenetic analysis of *E. faecium* isolates (n = 39) and to identify the carriage of antibiotic resistance genes. A long-read sequencing approach was adopted to identify structural variations in the vancomycin resistance region of a vancomycin-variable *E. faecium* (VVE) and to uncover the resistance reversion mechanism in this isolate. RT-qPCR and RT-PCR were used to determine differences in the expression of *vanRS* and *vanHAX* among strains.

**Results:** The *E. faecium* strains isolated in the hospital haematology ward were extensively drug resistant and highly diverse. The notable expansion of ST262 among patients was the likely driver of a VRE outbreak. A VVE isolate was identified that could rapidly revert to a vancomycin-resistant state in the presence of vancomycin. Disruption of the *vanR* gene in this isolate by an IS*6*-family element impaired its response to vancomycin. However, when the isolate was evolved to vancomycin resistance, it could constitutively express the *vanHAX* genes at levels up to 36,000-fold greater than the parent isolate via co-transcription with a ribosomal RNA operon.

**Conclusion:** In this study, we report a VVE isolate that was isolated during a VRE outbreak. This strain was capable of rapidly reverting to a resistant phenotype through a novel mechanism involving integration of *vanHAX* downstream of a ribosomal RNA operon. During VRE outbreaks, attention should be paid to contemporaneous vancomycin-susceptible strains as these may carry silent vancomycin resistance genes that can be activated through genomic rearrangements upon exposure to vancomycin.

## Introduction

*Enterococcus faecium* is a Gram-positive bacterium that is a commensal of the human gastrointestinal tract^1^. However, it is also an opportunistic pathogen that can cause bacteriaemia, endocarditis and urinary tract infections in immunocompromised hosts^2^. Genomic studies have revealed that the vast majority of clinical infections are caused by a phylogenetically defined cluster of *E. faecium* strains, which was termed clade A1^1^. *E. faecium* infections are difficult to treat as they are often resistant to aminoglycoside, fluoroquinolone, β-lactam, and glycopeptide drugs^3^.

Vancomycin is a bactericidal glycopeptide antibiotic that targets peptidoglycan of the bacterial cell wall^4^. Resistance to vancomycin is conferred by clusters of genes which replace the terminal D-alanyl D-alanine motif of the lipid II stem peptide with a D-alanyl D-lactate or D-alanyl D-serine motif, thereby greatly reducing the binding affinity of vancomycin^5^. There are currently 9 known gene clusters that confer resistance to vancomycin in *E. faecium*, but the *vanA* and *vanB*-type clusters are the most prevalent^6^. Vancomycin resistance gene clusters are generally carried on integrative and mobilizable elements of which Tn*1546* and Tn*1549* encode *vanA*- and *vanB-*type resistance, respectively^7,8^.

An increasing number of *E. faecium* strains are being identified that contain the gene clusters required for vancomycin resistance but are phenotypically susceptible^9–11^. These strains are known as vancomycin-variable *E. faecium* (VVE)^12^. The mechanisms which lead to the susceptibility of these isolates are varied. Full or partial deletion of genes within the vancomycin resistance gene cluster is common, including deletion of the regulatory genes *vanR*-*vanS*, the D-alanyl D-alanine dipeptidase gene *vanX*, or partial deletion of the D-alanyl D-alanine ligase gene *vanA*^12–16^. As well as gene deletions, integration of insertion sequence (IS) elements into the promoter regions of vancomycin resistance genes has also led to the evolution of VVE strains^17,18^. Vancomycin-variable isolates are of particular concern in the treatment of patients as some of these isolates can rapidly revert to the resistant phenotype under vancomycin selection, which may, in turn, lead to treatment failure.

Here we investigate an outbreak of vancomycin resistant *Enterococcus faecium* in a haematology ward within a UK hospital. Within the outbreak we identified a vancomycin-variable isolate that was able to rapidly revert to a vancomycin-resistant phenotype under low-level vancomycin selection and we uncovered both the cause of its susceptibility and the mechanism by which it could revert to a vancomycin resistant phenotype.

## Materials and methods

### Collection and isolation of *Enterococcus faecium*

*Enterococcus faecium* strains were isolated from a haematology ward in a hospital in Birmingham (United Kingdom) over a two-year period (2016-2017) of increased vancomycin resistant *E. faecium* (VRE) bacteraemia. 39 isolates were collected in total from 24 patients by blood culture and rectal screening. The blood culture samples were taken from febrile patients, while rectal screening samples were collected from all patients on the ward. Only vancomycin-resistant rectal screening isolates from patients with VSE bacteraemia were included in this study. Bacteria were initially isolated on Columbia CNA agar (Oxoid) plates and were confirmed as *Enterococcus faecium* by MALDI-TOF (Bruker). The *vanA*^+^ isolate *E. faecium* E8202 was used as a control for gene expression in Tn*1546*^19^.

### Short- and long-read sequencing

DNA extraction and whole genome shotgun sequencing (WGS) using Illumina technology was carried out by MicrobesNG (http://www.microbesng.com). Isolates were lysed by suspending in TE buffer (Invitrogen) containing 0.1 mg/ml lysozyme (Thermo Scientific) and 0.1 mg/ml RNase A (ITW Reagents), the suspension was incubated at 37°C for 25 mins. Proteinase K (VWR Chemicals) and SDS (Sigma Aldrich) were added to a final concentration of 0.1 mg/ml and 0.5% v/v respectively and incubated for a further 5 mins at 65°C. DNA was purified using an equal volume of SPRI beads and resuspended in EB buffer (Qiagen). DNA libraries were prepared using the Nextera XT Library Prep Kit (Illumina) and pooled libraries were sequenced on an Illumina HiSeq instrument using a 250 bp paired-end protocol.

High molecular weight DNA was extracted from isolate OI25 and its revertants using the Monarch® HMW DNA Extraction Kit for Tissue (New England Biolabs) according to the manufacturer’s protocol with the addition of 50 μg/ml lysozyme (Sigma Aldrich) to weaken the cell wall during the lysis step. The DNA libraries were prepared using the ligation sequencing kit SQK-LSK109 (Oxford Nanopore Technologies) and sequenced on the MinION platform (Oxford Nanopore Technologies) using a R9.4.1 flowcell (Oxford Nanopore Technologies).

Raw sequencing reads have been deposited in the European Nucleotide Archive under accession number PRJEB57409.

### DNA assembly

Adapters were removed from the short-read data and quality trimmed using fastp v.0.20.1^20^. Reads less than 50 bp were discarded and a sliding window quality cut-off of 15 was used. The short-read data was then assembled using shovill v.1.0.4 (https://github.com/tseemann/shovill) using the default parameters. Hybrid assemblies were created by Unicycler v.0.4.8^21^ using both short and long reads, Unicycler was run using the default parameters. Both the short-read and hybrid assemblies were annotated using PROKKA v.1.14.6^22^.

Hybrid assemblies of the VVE strain OI25 and its revertants have been deposited in Genbank under accession numbers GCA_947511065.1, GCA_947511075.1 and GCA_947510805.1.

### Phylogenetic analysis

A core genome alignment was created with Panaroo v.1.2.2^23^ using –clean-mode strict. Phylogenetic trees were created from the core genome alignments using RAxML v.8.1.15^24^ implementing the GTRGAMMA substitution model with 100 bootstraps. Recombination was removed from the trees using ClonalFrameML v.1.12^25^. Trees were midpoint rooted and visualised using iTOL v.5^26^. Isolates were typed with PubMLST^27^ using mlst v.2.18.0 (Seemann T, mlst, Github https://github.com/tseemann/mlst).

### Identification of antibiotic resistance determinants in *E. faecium* genomes

Antibiotic resistance genes were identified in the *E. faecium* isolates by querying the short-read assemblies against the ResFinder database^28^ using ABRicate v.0.9.8 (Seemann T, Abricate, Github https://github.com/tseemann/abricate). A minimum identity and coverage cut-off of 95% and 50% respectively was used to determine that the antibiotic resistance genes were present.

### Antibiotic susceptibility testing

All outbreak isolates were tested for their antibiotic susceptibility using the VITEK2 system (Biomérieux). A subset of the isolates was also tested using the broth microdilution method^29^ and interpreted with the EUCAST breakpoints. Assays were carried out in biological triplicate and the mode of the minimum inhibitory concentration was recorded. *E. faecium* E745 was used as a positive control in all assays^30^.

### Reversion of VVE from a vancomycin susceptible to vancomycin resistant phenotype

A colony of isolate OI25 was inoculated into 5 ml of Brain Heart Infusion (BHI) broth (VWR) and grown at 37°C for 16 hours with shaking (200 rpm). The culture was then diluted 1:100 into 5 ml of BHI broth containing 8 μg/ml vancomycin. The culture was grown at 37°C (200 rpm) and observed every 24 hours for growth. When growth was observed, the culture was diluted 10^6^-fold and 100 μl was spread onto BHI agar plates containing 8 μg/ml vancomycin. Two colonies were picked from the plate and stored for further analysis.

### Reverse transcription-quantitative polymerase chain reaction (RT-qPCR) and Reverse transcription polymerase chain reaction (RT-PCR)

RNA was extracted from cells collected in mid-log phase (OD_600_ = 0.5) and cells that had been exposed to 8 μg/ml vancomycin at mid-log phase for one hour, using the Monarch® Total RNA Miniprep Kit (New England Biolabs). Residual DNA was removed by treating the RNA with TURBO DNase™ (Invitrogen). cDNA was synthesised from the total RNA using the Maxima First Strand cDNA Synthesis Kit for RT-qPCR (Thermo Scientific). qPCR was carried out using PrimeTime® Gene Expression Master Mix (2X) (Integrated DNA Technologies (IDT)) and PrimeTime® qPCR Assays (20X) (IDT), which contained the forward primer, reverse primer and probe, for the *vanRS* and *vanHAX* operons, and the *tufA* housekeeping control gene (Table S1). The qPCR reaction was performed in a QuantStudio 1 Real-Time PCR system (Applied Biosystems™) with the following program: 95°C for 3 mins, followed by 40 cycles of 95°C for 15 s and 60°C for 1 min. Fold expression was calculated using the Livak method relative to the internal control gene *tufA*^31^.

The cDNA of isolate OI25rev2 was also used to perform RT-PCR assays across the rRNA-*vanHAX* junction. RT-PCR reactions were carried out using DreamTaq 2x Mastermix (Thermo Fisher Scientific) and forward and reverse primers that bridged between the 23S rRNA gene and the *vanH, vanA* and *vanX* genes (Table S2). The reactions were performed in a Mastercycler Pro Thermal Cycler (Eppendorf) with the following program: 95°C for 3 mins, followed by 30 cycles of 95°C for 30 s, 50°C for 30 s and 72°C for 2 mins, followed by a final incubation at 72°C for 10 mins. A reaction with a sample from which reverse transcriptase was omitted was used to control for residual DNA.

### Terminator analysis

The rho-independent terminator of the ribosomal RNA gene operon was identified in isolate OI25 by analysing 100 nucleotides downstream of the 5S rRNA gene stop codon via RNAfold Web Server^32^. The output was then manually inspected to identify the typical A-tail, loop, T-tail structure of a rho-independent terminator^33^.

### Fitness evaluation

Bacterial fitness was evaluated by comparing the maximum growth rate (μ_max_; h^-1^) and maximum growth (maximum A_600_) of the revertant isolates compared to isolate OI25. Bacterial cultures were grown for 16 hours at 37°C in BHI broth, diluted 1:1000 in BHI broth and added to a clear flat-bottom 96-well plate. Wells were included that contained only BHI broth to control for changes in A_600_ not caused by bacterial growth. The 96-well plate was incubated at 37°C with agitation (240 rpm) for 16 hours, absorbance measurements (600 nm) were taken at 10-minute intervals using a Spark microplate reader (TECAN). The experiment was carried out in biological and technical triplicates.

Maximum growth rate and maximum growth were determined using the R package Growthcurver v.0.3.1^34^.

### Statistical analyses

Tests for determining statistical significance were performed as described in the text and implemented in GraphPad Prism v.9.4.1.

## Results

### Genome sequence analysis revealed a multi-clonal, nosocomial VRE outbreak

A total of 39 isolates were collected in this study from 24 patients. 26 of the isolates were phenotypically resistant to vancomycin and 13 were phenotypically susceptible (Table S3). Thirty-four of the isolates were from blood culture samples and five were isolated from rectal screening swabs of patients.

Phylogenetic analysis of the clinical *E. faecium* isolates uncovered a complex population of isolates belonging to clade A1 (Figure S1). Eight different sequence types (ST262, ST80, ST1478, ST780, ST117, ST203, ST412 and ST787) were isolated on the ward during the period of the outbreak. A dominant ST262 clone that was present in 13 patients was the likely driver of the outbreak within the haematology ward. While all isolates could be assigned to clade A1, they were distinct from the clade A1 reference isolates (Figure 1). The outbreak isolates contained a large repertoire of antibiotic resistance genes. Aminoglycoside resistance was common among the isolates, with isolates carrying between two and five aminoglycoside resistance genes. All outbreak isolates carried the *E. faecium* specific *aac(6’)-Ii* gene and 33 of the 39 isolates carried the *aac(6’)-aph(2’’)* gene^35^. Erythromycin resistance genes were also found in all outbreak isolates: *erm(B)* was the most common macrolide resistance gene and was found in 33 of the isolates. Tetracycline resistance genes were found in 29 isolates, including *tet(L)* and four different alleles of *tet(M)*. Vancomycin resistance was widespread in the isolates with 25 out of 39 isolates carrying vancomycin resistance genes, all of which were the *vanA*-type. It was noted that isolate OI25 was phenotypically susceptible to vancomycin but carried the *vanHAX* genes necessary to confer phenotypic resistance, which suggested that it was a vancomycin-variable *Enterococcus faecium* (VVE) isolate. To confirm the result of the VITEK 2 susceptibility testing, *E. faecium* isolate OI25 was subjected to a broth microdilution MIC against vancomycin. Isolate OI25 had a vancomycin MIC of 1 μg/ml, which is below the EUCAST clinical breakpoint of 4 μg/ml, confirming that this isolate was indeed susceptible to vancomycin despite carrying the genes required for phenotypic resistance to vancomycin.

**Figure 1:**
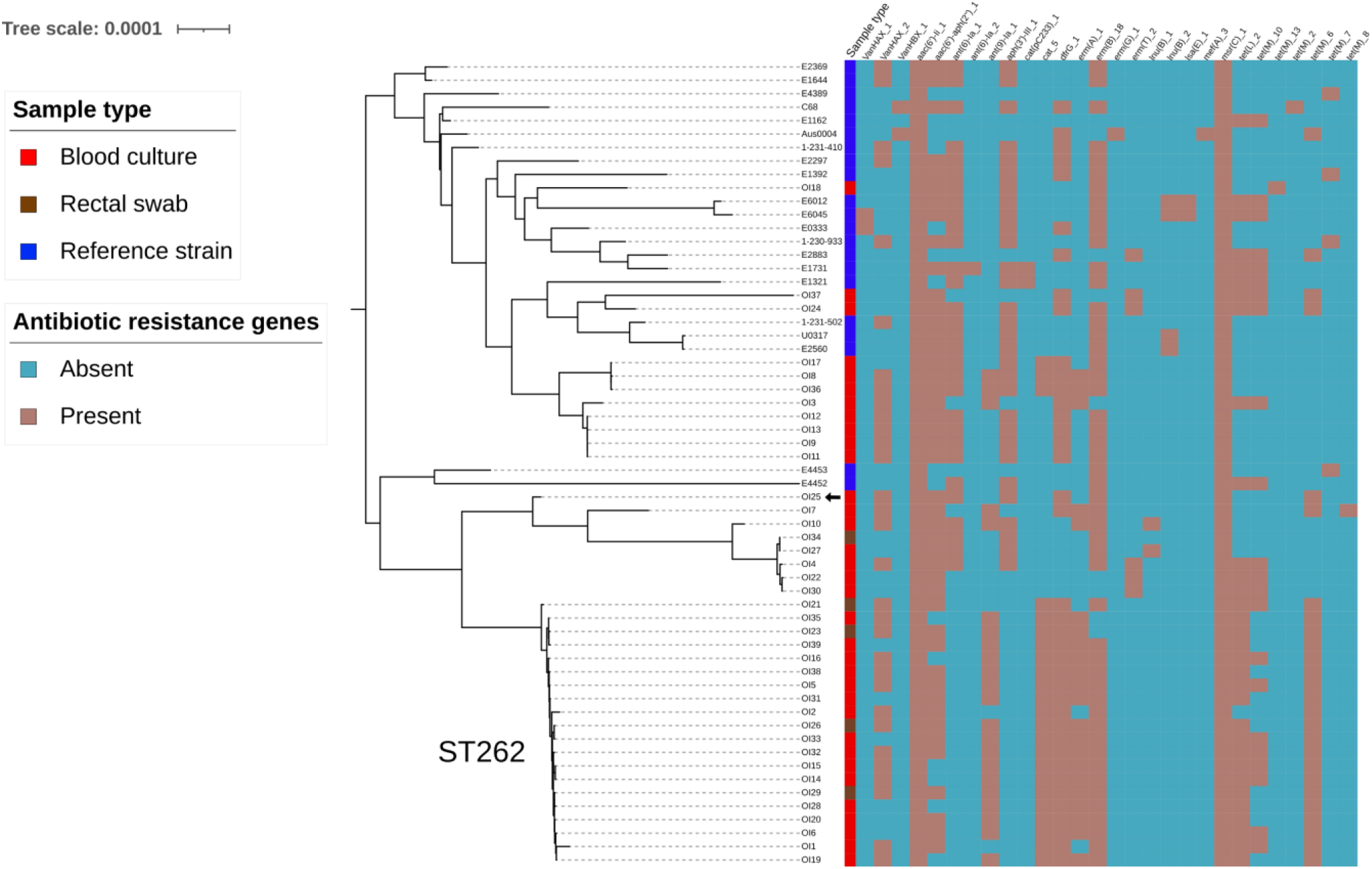
Maximum likelihood core genome phylogenetic tree of the clinical *E. faecium* isolates and representative clade A1 isolates. Metadata includes the sample type (Blood culture, Rectal swab or Reference strain) and the presence or absence of antibiotic resistance genes. The scale bar indicates the number of substitutions per site. The arrow indicates the VVE isolate OI25.

### OI25 had an impaired transcriptional response to vancomycin

RT-qPCR analysis was used to compare the transcriptional response of the vancomycin resistance operons *vanRS* and *vanHAX* in isolate OI25 to the vancomycin-resistant isolate E8202 when exposed to 8 μg/ml vancomycin (Figure 2). Expression of the *vanHAX* operon increased 310-fold in isolate E8202 when exposed to vancomycin but increased only 16-fold in isolate OI25. Similarly, upon exposure to vancomycin, expression of the *vanRS* genes increased 52-fold in the wildtype VRE isolate but only 5-fold in isolate OI25. This demonstrated that the susceptibility of isolate OI25 to vancomycin was due to its reduced ability to increase gene expression of the *vanHAX* operon in the presence of vancomycin.

**Figure 2:**
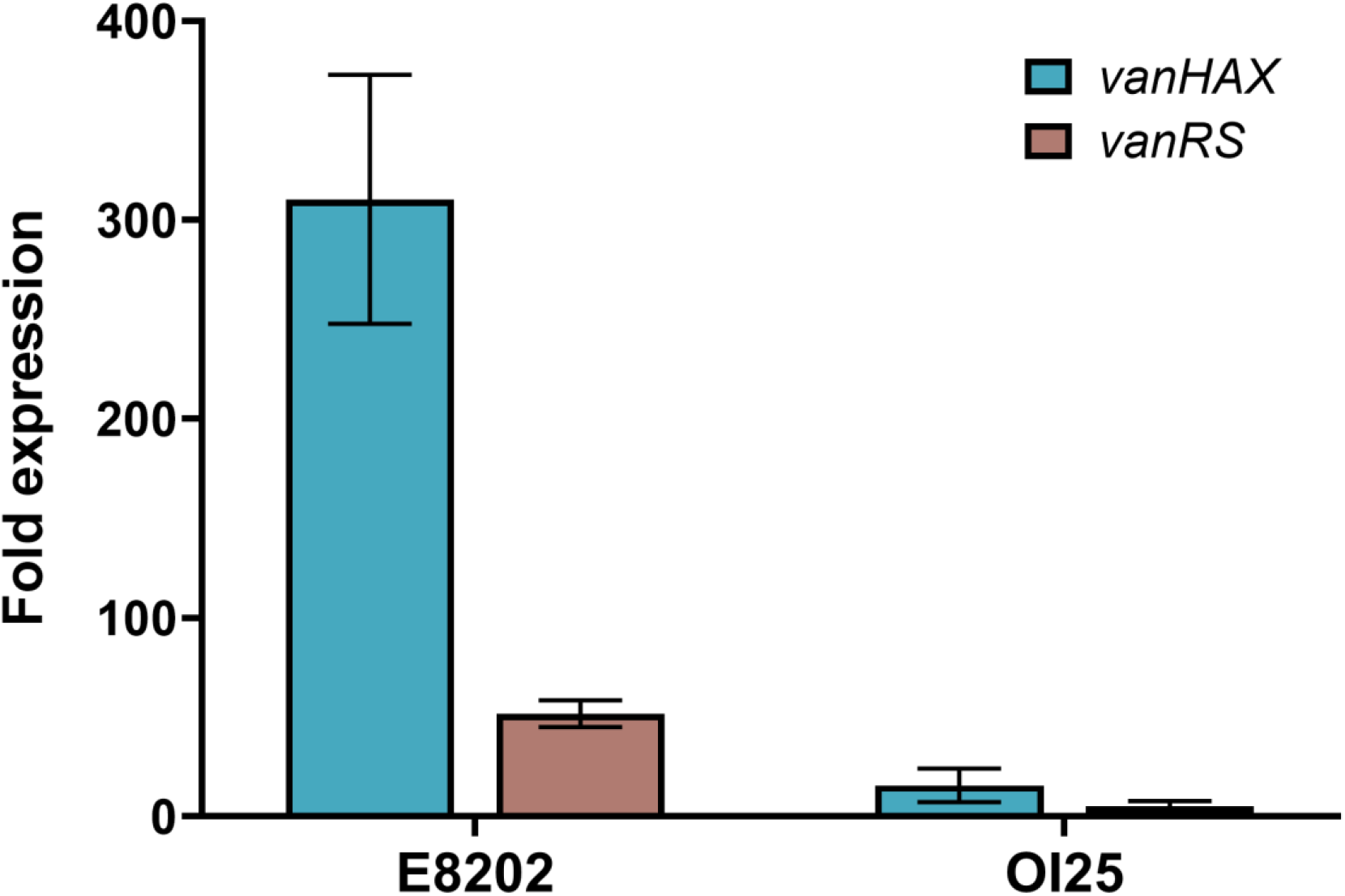
Expression of *vanHAX* and *vanRS* in *E. faecium* E8202 and the VVE isolate OI25. RT-qPCR analysis Expression of the vancomycin resistance gene operons *vanHAX* and *vanRS* of E8202 and the VVE isolate OI25 was determined by qRT-PCR analysis before and after exposure to 8 μg/ml vancomycin. Expression data was normalised to the internal control gene *tufA*. Experiments were carried out with biological triplicates and technical duplicates. Error bars represent standard deviation.

### An IS*6*-family element disrupted the *vanR* gene and its promoter in OI25

A genome assembly, incorporating both long- and short reads, of isolate OI25 was generated to analyse the vancomycin resistance region, in order to identify a possible mechanism that abolished vancomycin resistance in this isolate. Compared to the prototypical Tn*1546* transposon (GenBank: M97297.1), isolate OI25 had an insertion of an IS*L3*-family element between the *vanS* and *vanH* genes (Figure 3). However, this insertion did not occur within the previously characterised promoter region of *vanH* and thus did not disrupt the two VanR binding sites upstream of *vanH*^36^. OI25 also had an insertion of an IS*6*-family element within the promoter region and the first 50 bp of the *vanR* gene. It was likely that this inactivated *vanR*, thus preventing activation of the *vanHAX* genes in the presence of vancomycin, leading to the vancomycin-susceptible phenotype of OI25.

**Figure 3:**
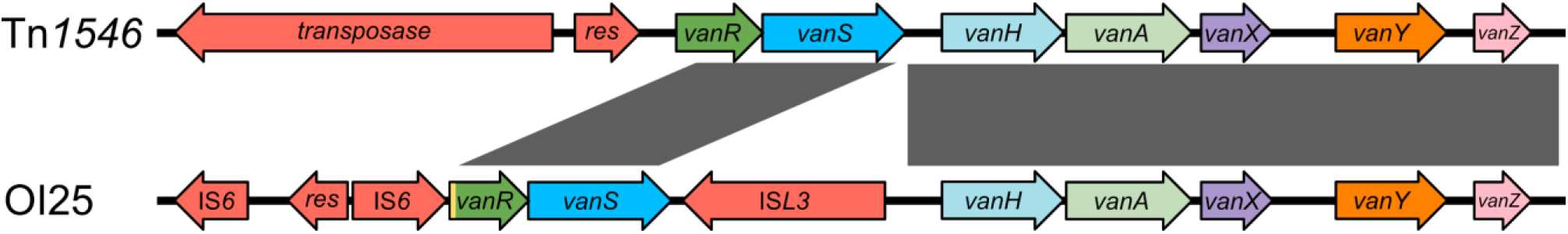
Alignment of the vancomycin resistance region of *E. faecium* isolate OI25 against the vancomycin resistance region of the prototypical Tn*1546* transposon. The Tn*1546* sequence was obtained from NCBI Genbank (accession number: M97297.1). Grey boxes represent regions which are identical between isolates. The yellow box represents the deletion in *vanR*.

### Rapid reversion to a high-level vancomycin resistant phenotype

Isolate OI25 was exposed to 8 μg/ml vancomycin to investigate whether it could revert to a vancomycin-resistant phenotype in the presence of a low concentration of vancomycin. Growth was observed within the OI25 culture after 48 hours. Two isolates taken from this culture had a vancomycin MIC of 512 μg/ml, thus showing that isolate OI25 could revert to a vancomycin resistant phenotype under vancomycin selection.

### Insertion of vancomycin resistance genes downstream of a ribosomal RNA operon led to a vancomycin-resistant phenotype

Complete genome assemblies of OI25 and its revertant isolates were generated by combining short- and long-read data, to identify a potential mechanism behind the reversion of isolate OI25 to a vancomycin-resistant phenotype. Both OI25 revertant isolates (OI25rev1 and OI25rev2) had similar genomic rearrangements compared to the parent isolate (Figure 4). The rearrangements led to the insertion of the vancomycin resistance genes into the chromosome, with the *vanHAX* operon becoming inserted immediately downstream of a ribosomal RNA operon, whereas the *vanRS* operon was inserted in such a way that it and its surrounding DNA remained unchanged. In isolate OI25rev2 the entire plasmid was integrated into the chromosome whereas in isolate OI25rev1 a 15,299-bp fragment of the plasmid DNA was integrated in the chromosome while a 21,107-bp plasmid remained. It could not be ascertained whether in isolate OI25rev1 the whole plasmid was integrated and then excised leaving the vancomycin resistance genes behind in the chromosome (Figure 4; green arrows) or whether the vancomycin resistance genes were excised and formed an intermediate mobile genetic element that was then integrated into the chromosome (Figure 4; purple arrows). In both isolates there was an 8-bp target site duplication (ACTAGAAA) surrounding the DNA inserted into the chromosome that is consistent with the action of an IS element.

**Figure 4:**
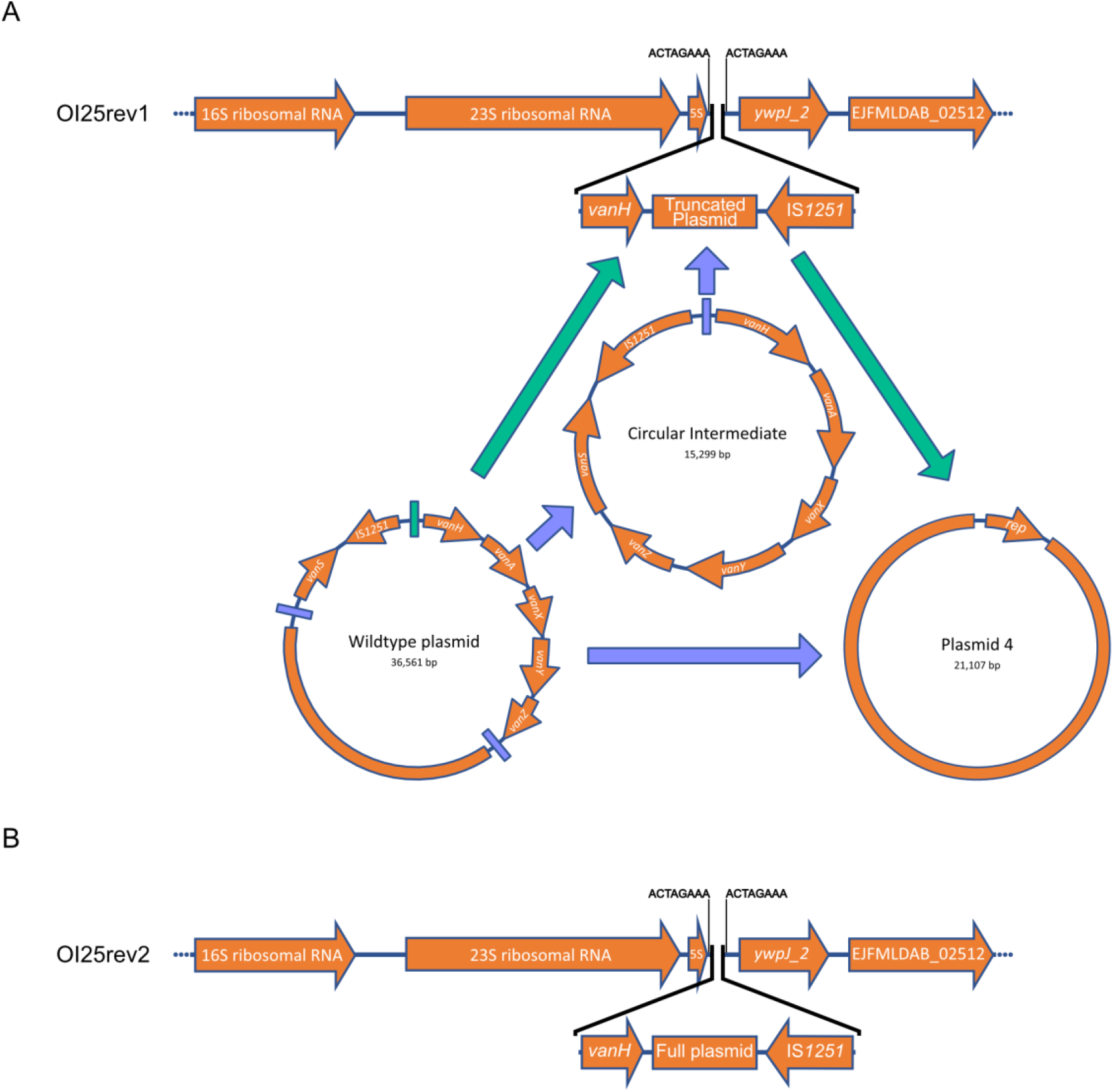
Mechanisms of VVE reversion to vancomycin resistance. A. Insertion of the vancomycin resistance plasmid into the chromosome of OI25rev1 and the possible intermediate stages in the insertion. B. Insertion of the vancomycin resistance genes into the chromosome of OI25rev2.

### Substantial, constitutive upregulation of *vanHAX* expression in revertant isolates

As the insertion of the plasmid DNA into the chromosome did not restore the *vanR* gene it was hypothesised that the *vanHAX* genes were instead being constitutively expressed. To determine whether the expression of the *vanHAX* operon had changed at this new locus, RT-qPCR was used to compare the expression of the *vanHAX* and *vanRS* operons in the revertant isolates compared to OI25. Although the insertions that occurred in both revertant isolates were different, the change in expression of the *vanHAX* and *vanRS* operons was similar. In the absence of vancomycin, the expression of the *vanHAX* operon in the two revertant strains OI25rev1 and OI25rev2 was on average (± standard deviation) 2.7 × 10^4^ ± 1.1 × 10^4^-fold and 3.6 × 10^4^ ± 1.3 × 10^4^-fold greater than in OI25. The expression of the *vanRS* operon was also 39.4 ± 22.8-fold (OI25rev1) and 34.2 ± 16.3-fold (OI25rev2) higher in the revertants, compared to OI25, despite the continued disruption of the *vanR* gene. This demonstrated that the revertant isolates were expressing the *vanHAX* genes needed to confer resistance to vancomycin, even in the absence of vancomycin.

The genomic insertion site was inspected in both the revertant and parent isolates to determine a mechanism behind the constitutive expression of the *vanHAX* genes. In the parental OI25 strain, a putative rho-independent terminator of the ribosomal RNA operon was uncovered (Figure S2). The chromosomal insertion of plasmid DNA in isolates OI25rev1 and OI25rev2 occurred 27 bp downstream of the 5S rRNA gene stop codon. This insertion occurred approximately halfway through the putative rho-independent terminator leading to the disruption of its secondary structure. It was hypothesised that disruption of the rho-independent terminator could lead to the co-transcription of the ribosomal RNA genes and the *vanHAX* genes.

To determine whether the *vanHAX* operon was co-transcribed with the upstream ribosomal RNA gene operon, RNA was reverse transcribed from isolates OI25rev1 and OI25rev2 and PCR was performed across the rRNA - *vanHAX* operon junction. Three PCR reactions were performed on the cDNA each of which spanned from the 23S ribosomal RNA gene into the *vanH, vanA* and *vanX* genes (Figure 5). PCR amplicons of the expected lengths were present for all three gene which confirmed that the *vanHAX* genes were indeed co-transcribed with the ribosomal RNA genes.

**Figure 5:**
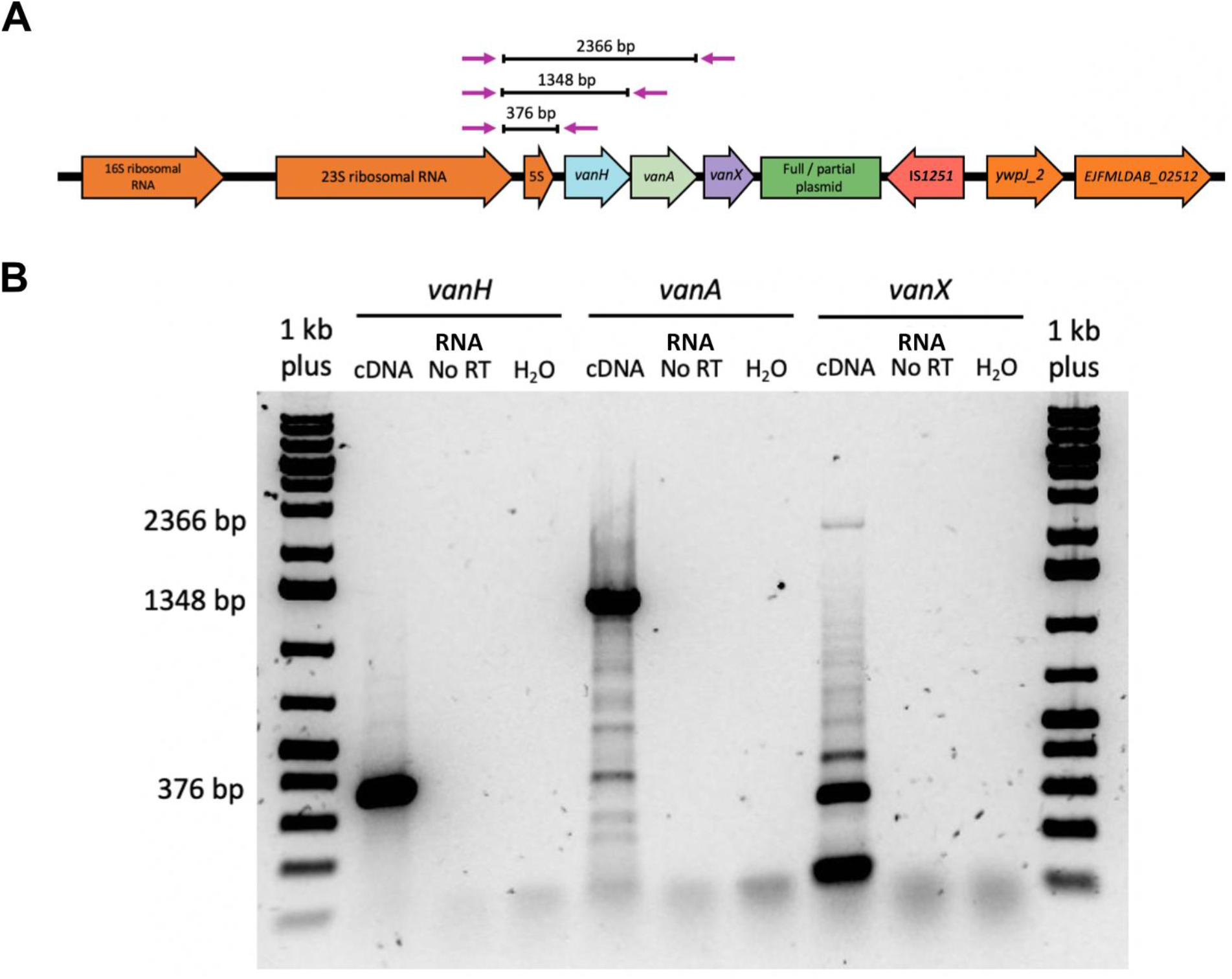
RT-PCR on the rRNA-*vanHAX* junction in OI25rev2. **A**. Schematic showing the expected amplicon sizes. **B**. 1% agarose gel showing the amplicons with the expected products sizes from panel **A** indicated for the RT-PCR reactions between the 23S rRNA gene and *vanH* (376 bp), *vanA* (1348 bp) and *vanX* (2366 bp). Ladder: GeneRuler 1kb Plus (Thermo Scientific). RT = Reverse Transcriptase.

### Vancomycin-resistant revertant isolates do not show a growth defect

It was hypothesised that co-transcription of the vancomycin resistance genes with the ribosomal RNA genes would impose a high fitness cost in the revertant isolates. However, when the wildtype isolate OI25 and its revertants were grown in the absence of vancomycin selection (Figure S3), μ_max_ of isolate OI25 (1.7 h^-1^) was not significantly different to that of OI25rev1(1.7 h^-1^, Kruskal-Wallis, *P* > 0.99) or OI25rev2 (1.8 h^-1^, Kruskal-Wallis, *P* = 0.11). Similarly, the maximum growth reached by OI25rev1 (A_600_ 0.29) and OI25rev2 (A_600_ 0.25) was lower, but not significantly different, from that of OI25 (A_600_ 0.36; Kruskal-Wallis versus OI25rev1 *P* = 0.22 and versus OI25rev2 *P* = 0.08). Despite the vancomycin resistance genes being transcribed at a high level in the revertant isolates, this did not impose a significant fitness cost.

## Discussion

The present study aimed to investigate a VRE outbreak in a haematology ward. The isolates in this study belonged to 8 different sequence types within the hospital-associated clade A1^1^. The major clone driving the outbreak belonged to ST262, with the presence of highly related ST262 isolates in 13 different patients suggesting spread within the ward. ST262 has previously been associated with the hospital environment in the UK and Europe but has not thus far been identified as a prominent driver of a VRE outbreak^37–39^. Other isolates belonged to ST80 which has been linked to VRE outbreaks in Ireland and Sweden^40,41^.

An isolate (OI25) belonging to ST787 was identified that was genotypically resistant to vancomycin but phenotypically susceptible. Long-read sequencing uncovered multiple IS element insertions into the vancomycin resistance regions compared to the wildtype transposon Tn*1546*^7^. An IS*L3* family element was inserted between the *vanS* and *vanH* genes. This insertion likely did not contribute to the susceptibility of the isolate as it occurred outside of the promoter region and an identical insertion has been found in other isolates which maintain a resistant phenotype^42^. There was also a further insertion of an IS*1216* element into the promoter region and first 50 bp of the *vanR* gene. This insertion was unique among global isolates, but a similar vancomycin-variable *E. faecium* isolate has been described, which also contained an insertion of an IS*1216* family element that deleted the first 55 bp of the *vanR* gene^17^. As the insertion of the IS*1216* element occurred within the *vanR* gene and its promoter region it was likely that isolate OI25 could not respond to vancomycin which was subsequently confirmed by RT-qPCR.

Although vancomycin-variable enterococci are phenotypically susceptible to vancomycin, some isolates can revert to a resistant phenotype under antibiotic selection. Several mechanisms have been uncovered including the excision of IS elements leading to the formation of constitutive promoters^17^, gene duplication events^43^ and the acquisition of plasmids containing vancomycin resistance genes^15^. Exposure of isolate OI25 to 8 μg/ml vancomycin led to a rapid reversion of the isolate to high-level vancomycin resistance. Long-read read sequencing of two revertant isolates uncovered that the vancomycin resistance genes *vanH, vanA* and *vanX* had become inserted into the chromosome directly downstream of a ribosomal RNA operon. This insertion caused a disruption of the rho-independent terminator of the operon and led to the co-transcription of the vancomycin resistance genes in a constitutive manner. The native high-level expression of the ribosomal RNA genes led to a significant upregulation in the *vanHAX* genes^44^. The presence of an 8 bp target site duplication and an IS*1251* family element at the 3’ end of the inserted DNA suggested an IS mediated rearrangement of the DNA through a currently uncharacterised mechanism^45^.

Our findings highlight the diversity of mechanisms that enable VVE isolates to revert to their resistant state. While vancomycin-variable *E. faecium* typically make up a small percentage of the *E. faecium* strains isolated within the clinical environment, they have in places become the dominant clone^10^. As VVE isolates become more common in the hospital environment it may be necessary for the successful treatment of these infections to include whole genome long-read sequencing in routine pathogen diagnostics to rapidly identify strains that are phenotypically susceptible to vancomycin but can potentially revert to high-level vancomycin resistance.

## Acknowledgements

We would like to thank the laboratory staff at UKHSA for the collection of the *E. faecium* isolates and MicrobesNG for genome sequencing of *E. faecium* isolates.

## Author contributions

W.V.S. and A.H. conceived this study. A.H. and K.H. collected the bacterial isolates. R.S.M., A.E.S., M.P., S.J.D and W.V.S. prepared the samples for sequencing. R.S.M., A.E.S and S.J.D. analysed the data. R.S.M. and W.V.S. wrote the manuscript with input from all authors.

## Ethical statement

This study did not require ethical approval as it was part of a hospital infection control investigation into a local outbreak.

