## Supplementary material for "A novel resistance reversion mechanism in a vancomycin-variable *Enterococcus faecium* strain": Figure S1

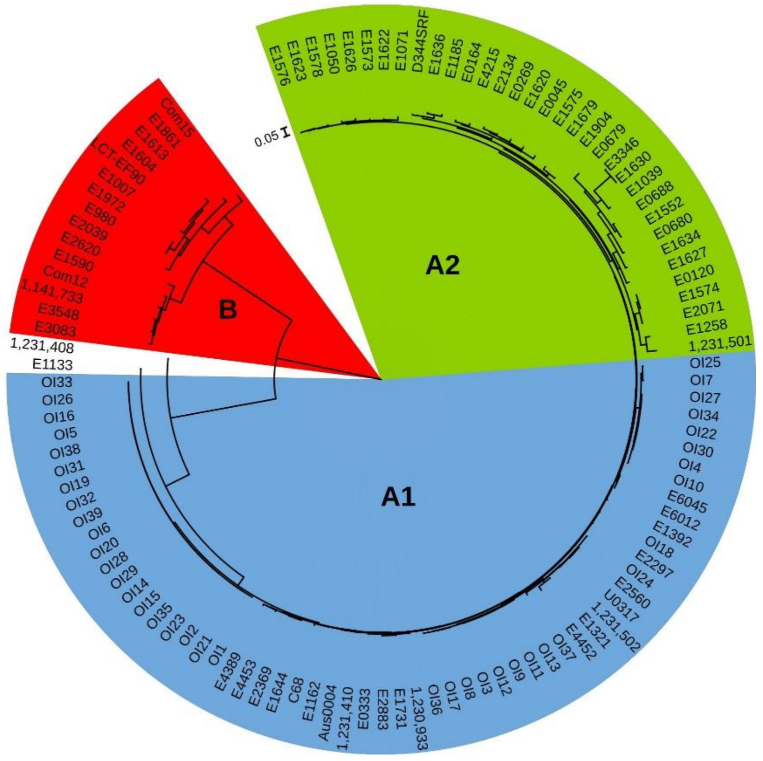


**Figure S1: Maximum likelihood core genome phylogenetic tree of the outbreak isolates with representative *E. faecium* isolates from clades A1, A2 and B.** The major clades have been coloured, clade A1 - blue, clade A2 - green and clade B - red. The scale bar indicates the number of substitutions per site.
