## Supplementary material for "A novel resistance reversion mechanism in a vancomycin-variable *Enterococcus faecium* strain": Figure S2

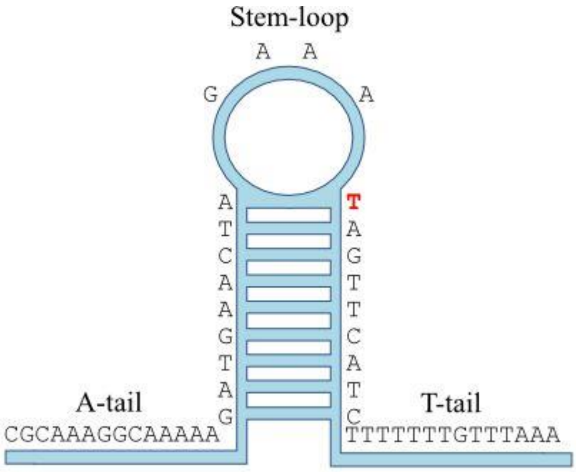


**Figure S2: The predicted rho-independent terminator of the 5S rRNA gene.** The terminator was found within 50 bp of the predicted 5S rRNA gene end. The insertion site of the vancomycin resistance plasmid is highlighted in red.
