## Supplementary material for "A novel resistance reversion mechanism in a vancomycin-variable *Enterococcus faecium* strain": Figure S3

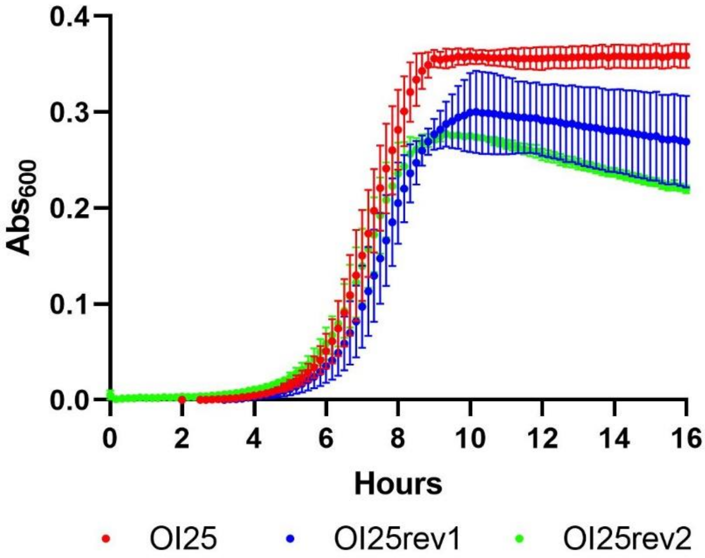


**Figure S3: Bacterial growth curves of isolate OI25 (red), OI25rev1 (blue) and OI25rev2 (green) in BHI broth.** Error bars represent standard deviation. The experiment was carried out in biological and technical triplicates.
