## Supplementary material for "A novel resistance reversion mechanism in a vancomycin-variable *Enterococcus faecium* strain": Table S1

**Table S1: RT-qPCR primers to measure expression of *vanHAX* and *vanRS* operons.**

| Primer name | Sequence (5' -> 3') | Description |
|---|---|---|
| tufA_fwd | GGTGACGATGTTCCTGTAGTT | PrimeTime qPCR Probe assay targeting the *tufA* gene in *E. faecium*. |
| tufA_probe | TGAAAGCTCTAGAAGGCGACGCTT | |
| tufa_reverse | CGTTCTGGAGTTGGGATGTATT | |
| vanHAX_fwd | ATATAAAGCGCTCGGCTGTAG | PrimeTime qPCR Probe assay targeting the *vanHAX* operon in *E.faecium*. |
| vanHAX_probe | TAACGGCCGCATTGTACTGAACGA | |
| vanHAX_rev | TGAAACCGGGCAGAGTATTG | |
| vanRS_fwd | AAGCTGGCCGAACAAAGA | PrimeTime qPCR Probe assay targeting the *vanRS* operon in *E.faecium*. |
| vanRS_probe | ACGTTGTTATGTACTTGGCGCACG | |
| vanRS_rev | CGTCAAGCAGGCTCAAATAAC | |
