## Supplementary material for "A novel resistance reversion mechanism in a vancomycin-variable *Enterococcus faecium* strain": Table S2

**Table S2: RT-PCR primers to test for co-transcription of rRNA genes and *vanHAX* genes.**

| Prime name | Sequence (5' -> 3') | Description |
|---|---|---|
| Junc_fwd | CGGACTGATACTAATCGATCG | Primer used to check for junction between rRNA and *vanHAX* genes in cDNA. |
| Junc_vanH_rev | GGAATGCATCTGCCTCATC | Primer used to check for junction between rRNA and *vanH* gene in cDNA. |
| Junc_vanA_rev | AGATTTTACCGATACGTCATGC | Primer used to check for junction between rRNA and *vanA* gene in cDNA. |
| Junc_vanX_rev | CAACGAACACCGTGTACTAT | Primer used to check for junction between rRNA and *vanX* gene in cDNA. |
