## Supplementary material for "A novel resistance reversion mechanism in a vancomycin-variable *Enterococcus faecium* strain": Table S3

**Table S3. Metadata of strains collected in this study**

| Isolate name | Patient | Isolation Month | Sample type | MLST | Vancomycin phenotype | Vancomycin genotype |
|---|---|---|---|---|---|---|
| OI1 | 23 | Mar 2016 | Blood Culture | 262 | Resistant | Resistant |
| OI2 | 1 | Apr 2016 | Blood Culture | 262 | Resistant | Resistant |
| OI3 | 2 | May 2016 | Blood Culture | 1478 | Resistant | Resistant |
| OI4 | 3 | May 2016 | Blood Culture | 80 | Resistant | Resistant |
| OI5 | 4 | Jun 2016 | Blood Culture | 262 | Resistant | Resistant |
| OI6 | 6 | Feb 2017 | Blood Culture | 262 | Resistant | Susceptible |
| OI7 | 7 | Feb 2017 | Blood Culture | 80 | Resistant | Resistant |
| OI8 | 8 | Feb 2017 | Blood Culture | 780 | Resistant | Resistant |
| OI9 | 9 | Apr 2017 | Blood Culture | 1478 | Resistant | Resistant |
| OI10 | 10 | Apr 2017 | Blood Culture | 80 | Resistant | Resistant |
| OI11 | 11 | May 2017 | Blood Culture | 1478 | Resistant | Resistant |
| OI12 | 12 | May 2017 | Blood Culture | 1478 | Resistant | Resistant |
| OI13 | 13 | May 2017 | Blood Culture | 1478 | Resistant | Resistant |
| OI14 | 14 | May 2017 | Blood Culture | 262 | Resistant | Resistant |
| OI15 | 14 | Jun 2017 | Blood Culture | 262 | Resistant | Resistant |
| OI16 | 15 | Jun 2017 | Blood Culture | 262 | Resistant | Resistant |
| OI17 | 24 | Jul 2017 | Blood Culture | 780 | Susceptible | Susceptible |
| OI18 | 7 | Feb 2017 | Blood Culture | 117 | Susceptible | Susceptible |
| OI19 | 5 | Dec 2016 | Blood Culture | 262 | Resistant | Resistant |
| OI20 | 17 | Aug 2017 | Blood Culture | 262 | Susceptible | Susceptible |
| OI21 | 22 | Nov 2017 | Rectal Swab | 262 | Resistant | Resistant |
| OI22 | 18 | Nov 2017 | Blood Culture | 80 | Susceptible | Susceptible |
| OI23 | 18 | Nov 2017 | Rectal Swab | 262 | Resistant | Resistant |
| OI24 | 13 | May 2017 | Blood Culture | 412 | Susceptible | Susceptible |
| OI25 | 1 | Dec 2016 | Blood Culture | 787 | Susceptible | Resistant |
| OI26 | 19 | Oct 2017 | Rectal Swab | 262 | Resistant | Resistant |
| OI27 | 20 | Nov 2017 | Blood Culture | 80 | Susceptible | Susceptible |
| OI28 | 21 | Aug 2017 | Blood Culture | 262 | Susceptible | Susceptible |
| OI29 | 21 | Sep 2017 | Rectal Swab | 262 | Resistant | Resistant |
| OI30 | 22 | Aug 2017 | Blood Culture | 80 | Susceptible | Susceptible |
| OI31 | 4 | Jun 2016 | Blood Culture | 262 | Susceptible | Susceptible |
| OI32 | 6 | Feb 2017 | Blood Culture | 262 | Resistant | Resistant |
| OI33 | 19 | Oct 2017 | Blood Culture | 262 | Susceptible | Susceptible |
| OI34 | 20 | Nov 2017 | Rectal Swab | 80 | Resistant | Susceptible |
| OI35 | 16 | Jul 2017 | Blood Culture | 262 | Resistant | Resistant |
| OI36 | 17 | Aug 2017 | Blood Culture | 780 | Resistant | Resistant |
| OI37 | 12 | May 2017 | Blood Culture | 203 | Susceptible | Susceptible |
| OI38 | 4 | Jun 2016 | Blood Culture | 262 | Resistant | Resistant |
| OI39 | 6 | Feb 2017 | Blood Culture | 262 | Susceptible | Susceptible |
| OI25rev1 | NA | NA | Laboratory evolved | 787 | Resistant | Resistant |
| OI25rev2 | NA | NA | Laboratory evolved | 787 | Resistant | Resistant |
